# Genome editing of marsupial *RSX* reveals conserved and divergent principles in mammalian X-inactivation

**DOI:** 10.64898/2026.07.01.735830

**Authors:** Aurélien Courtois, Sergio Menchero, Sugako Ogushi, Wazeer Varsally, Sophie Wood, Fanny Decarpentrie, Riko Yoshimi, Aki Shiraishi, Kenichi Inoue, Takaya Abe, John L. VandeBerg, Daniel M. Snell, Hiroshi Kiyonari, James M.A. Turner

## Abstract

X-chromosome inactivation balances X-gene dosage between the sexes in eutherians and marsupials. The lncRNA *Xist* mediates X-inactivation in eutherians but is absent from marsupials. An evolutionarily unrelated lncRNA, *RSX*, has been identified in marsupials, but its role in X-inactivation is unresolved because genome editing in these mammals has only recently become feasible. Using CRISPR-Cas9 in the opossum, we show that *RSX* is required for the initiation of X-inactivation in marsupial embryos. *RSX* deletion later, in differentiated cells, causes modest X-gene de-repression, with repressive chromatin marks H3K27me3 and H3K9me3 retained. H2AK119ub is not enriched on the opossum inactive X, revealing divergent Polycomb-mediated regulation in therians. Our findings reveal stage-specific roles of *RSX* in marsupial X-inactivation and identify conserved and divergent features of X-dosage compensation in mammals.

## Introduction

The different sex chromosome content between male (XY) and female (XX) mammals generates an X-dosage imbalance that is resolved by X-inactivation. This epigenetic process silences one of the two X chromosomes in female cells, but the modes by which X-inactivation is achieved diverge in the two major clades of mammals (*1*). In eutherians, X-inactivation in the epiblast is random with respect to X chromosome parent of origin and triggered by the long non-coding RNA (lncRNA) *Xist* (*2–5*), which ultimately directs H2AK119ub and H3K27me3 to the future inactive X (*6–10*). For most, but not all X-genes, long-term silencing is independent of *Xist* and likely ensured by promoter DNA methylation (*11–18*). In marsupials, X-inactivation is imprinted, and the paternal X is always silenced (*19*). Repressive histone marks, such as H3K27me3, also coat the inactive X (*20–22*), but DNA methylation is not enriched at promoters of silenced genes (*23–27*), raising questions about how X-inactivation is maintained. More strikingly, *Xist* is not found in marsupials (*28*) and instead, *RSX*, another lncRNA with no homology, but with *Xist*-like properties, was discovered (*29*, *30*).

Determining whether *RSX* mediates X-inactivation has not been possible due to the myriad challenges of CRISPR-Cas9 genome editing in marsupial embryos. To date, only one study, targeting a protein-coding gene in the opossum *Monodelphis domestica*, has been reported (*31*). Here, we developed a pipeline to target *RSX.* By combining *in vitro* and *in vivo* genome editing in the opossum, together with transcriptomics and epigenomics, we uncovered convergent and divergent strategies in mammalian X-inactivation, and demonstrated the potential of genome editing to resolve longstanding questions in marsupial biology.

## Results

### *RSX* deletion causes biallelic expression of an X-gene in female opossum embryos

To assess the role of *RSX* in marsupial X-inactivation, we generated a loss-of-function deletion using CRISPR-Cas9. We combined two guide RNAs, one targeting upstream of the promoter and the other targeting the 5’ end of exon 1, which together generated a 1.7kb deletion that extinguished *RSX* expression in opossum female fibroblasts (**fig. S1A-C**).

In the opossum, *RSX* expression occurs simultaneously with embryonic genome activation (EGA) at embryonic day (E) 3.5 (8-cell stage) (*30*). To investigate its role in the initiation of X-inactivation, we targeted *RSX* in zygotes using pronuclear microinjection, transferred them into pseudopregnant surrogate females, and harvested putatively edited embryos at E5.5 (32-128 cell-stage) (**Fig. 1A**). Subsequently, we performed RNA FISH for *RSX* and *MSN*, a gene subject to X-inactivation (*29*). As a control, we targeted the autosomal coat colour gene *TYR* (tyrosinase), which has been successfully edited previously (*31*).

**Fig 1.**
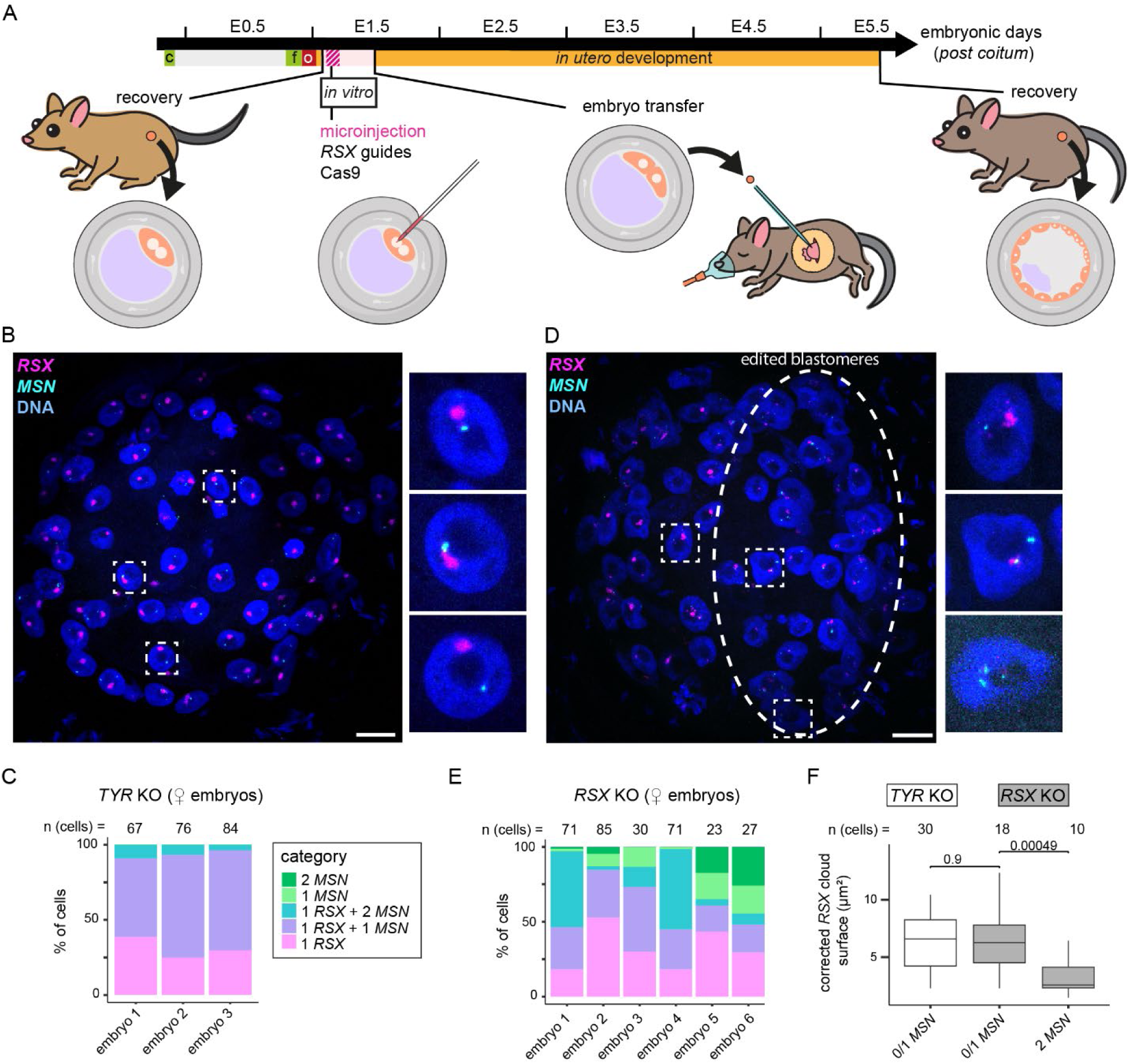
*RSX* deletion is associated with biallelic *MSN* expression in opossum E5.5 embryos. **A)** Schematic representation of the *RSX*-targeting strategy; (c) coitus, (f) fertilisation, (o) oviduct transport; tick marks indicate midnight between each day. **B)** Confocal Z-projection of female opossum embryo microinjected with guides targeting *TYR* (embryo 3) showing RNA FISH signal for *RSX* (magenta) and *MSN* (cyan). Scale bar 20µm. Insets show three representative cells. **C)** Bar plots showing percentage of nuclei in each category for the three *TYR* KO embryos. **n** indicates the number of cells showing a signal per embryo. Total cell numbers per embryo, including cells without any FISH signal, are 69, 76 and 86, respectively. **D)** Confocal Z-projection of female opossum embryo microinjected with guides targeting *RSX* (embryo 1). Scale bar 20µm. Dotted ellipse highlights area where fewer and smaller *RSX* clouds are visible. Insets show three representative cells. **E)** Bar plots representing the percentage of nuclei in each category for the six *RSX* KO embryos. **n** indicates the number of cells showing a signal per embryo. Total cell numbers per embryo, including cells without any FISH signal, are 104, 45, 81, 34, and 35 respectively. **F)** Boxplot of the *RSX* cloud maximal surface throughout the Z-stack for representative nuclei in three different embryos for the *TYR* and *RSX* KO. All *TYR* nuclei have 0 or 1 *MSN* dot, whereas *RSX* KO are split between 0 and 1 (monoallelic *MSN*) and 2 (biallelic *MSN*) *MSN* dots. **n** represents the number of total number of nuclei quantified across three embryos in each category.

Opossum embryos display developmental heterogeneity within litters. To ensure that we captured the stage when X-inactivation is complete, we first focused on microinjected embryos that had reached the 32-cell stage or later. In *TYR*-guide microinjected female embryos, blastomeres consistently exhibited an *RSX* cloud marking the inactive paternal X chromosome (Xi), and monoallelic expression of *MSN* from the active maternal X chromosome (Xa; **Fig. 1B** and **1C**). These findings are the expected outcome of successful X-inactivation, and mirrored published findings in non-microinjected opossum embryos (*30*).

In contrast, *RSX*-guide microinjected female embryos showed X-inactivation related phenotypes. All embryos were mosaics: while some blastomeres displayed *RSX* clouds and monoallelic *MSN* expression, many blastomeres showed diminished or absent *RSX* clouds, with biallelic *MSN* expression, suggestive of X-inactivation failure (**Fig. 1D** and **1E, fig. S1D** and **S1E**). Blastomeres with biallelic *MSN* exhibited reduced *RSX* cloud surface (measured in cross-sections) relative to those with monoallelic *MSN* (**Fig. 1F**), causally linking *RSX* to *MSN* silencing. *A priori*, the presence of smaller *RSX* clouds could be explained by delayed CRISPR-Cas9 genome editing or by smaller promoter deletions that diminish but do not extinguish *RSX* expression. Biallelic *MSN* expression was observed in up to 54% of blastomeres from *RSX*-guide microinjected female embryos, compared with 9% of *TYR*-microinjected controls. We conclude that *RSX* is necessary for opossum X-inactivation, as determined by RNA-FISH.

Our RNA-FISH analysis revealed that *RSX*-deletant embryos with ≥32 cells were mosaics. This finding could be explained by the extent and timing of CRISPR-Cas9 genome editing in zygotes. Alternatively, embryos with more efficient *RSX* deletion might exhibit developmental delay, manifested as a lower cell number. Supporting our second hypothesis, we recovered several embryos with highly efficient *RSX* deletion, including three with 100% *RSX* deletion efficiency, and all had fewer than 32 cells (**fig. S1F**). Thus, X-inactivation failure may cause counterselection early in opossum embryogenesis (also see later).

### *RSX* deletion causes an elevated X-to-autosome ratio

To gain a chromosome-wide view of how *RSX* deletion influences X-inactivation, we turned to single-cell multi-omics. We recovered microinjected embryos at E5.5 and performed combined single-cell whole-genome sequencing and transcriptome sequencing (scG&T-seq), thereby resolving CRISPR-induced *RSX* mutation signatures and their corresponding effects on X-gene expression. Sequencing of parental DNA, followed by genome-wide SNP calling, allowed us to discriminate maternal versus paternal alleles in our transcriptomic and genomic data. We recovered a total of 39 embryos from 9 females, from which we harvested 858 single cells for single-cell transcriptomics, 379 of which were retained after quality control. Cells were sexed using a previously described method (*30*), resulting in 228 female cells for further analysis (**fig. S2A**).

We determined the developmental stage of all female cells by integrating them into a reference dataset with non-injected E1.5-E5.5 opossum embryo cells ((*30*), and see **Materials and Methods**). Uniform manifold approximation and projection (UMAP; **Fig. 2A**) and Slingshot pseudotime analysis (**fig. S2B**) identified two predominant populations within our injected cells. One exhibited the expected E5.5-like signature, and the second, a delayed, E4.5-like signature (**Fig. 2B**). An additional smaller population exhibited an E1.5/2.5-like profile. These cells had failed to initiate EGA, as demonstrated by the enrichment of maternal and oocyte-related transcripts (**fig. S2C** and **table S1**), and were therefore not considered further.

**Fig 2.**
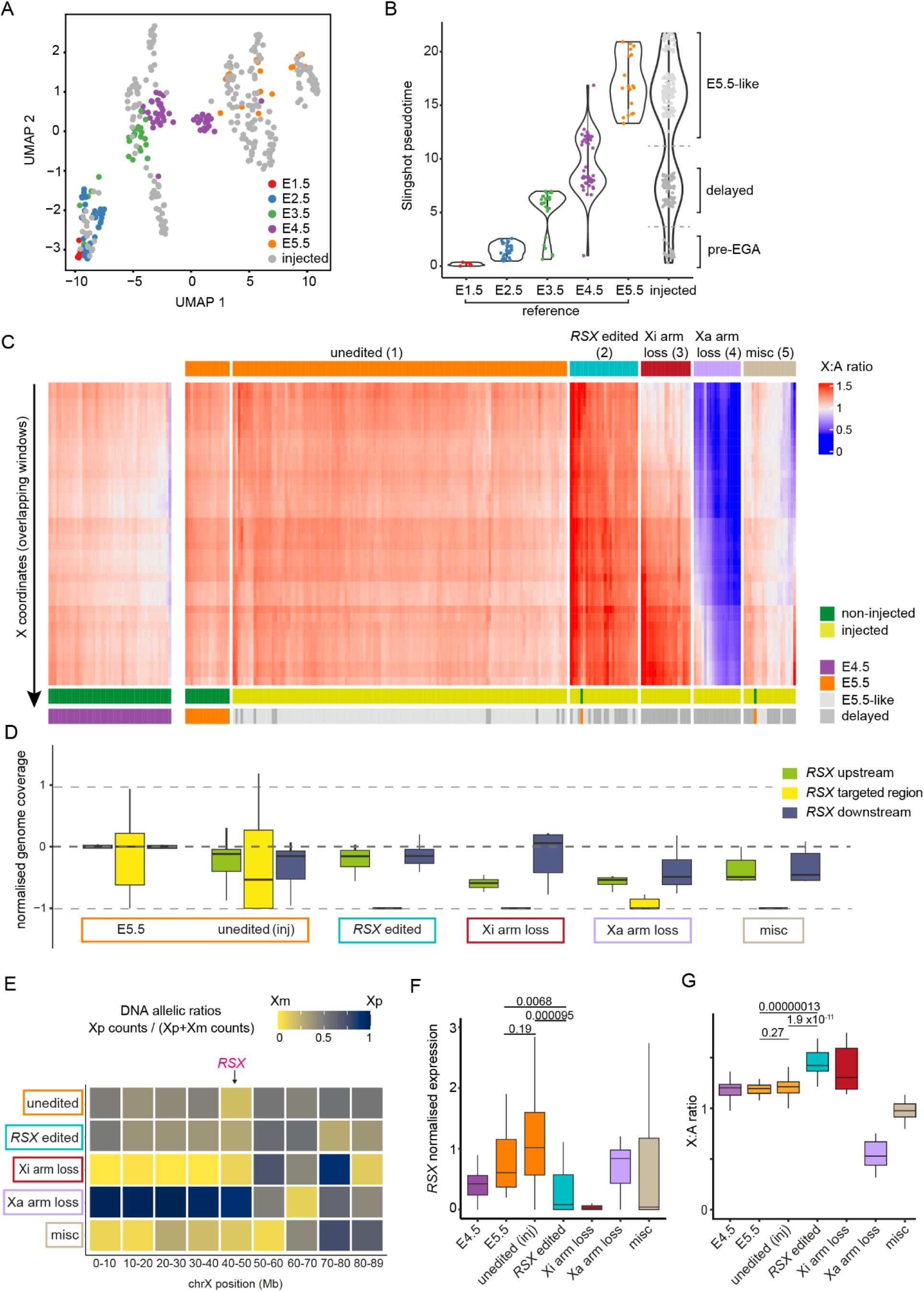
*RSX* deletion perturbs the initiation of X-inactivation. **A)** UMAP representation of single cells from all the XX E5.5 injected and from E4.5 and E5.5 non-injected embryos, as well as single cells from E1.5, 2.5 and 3.5 non-injected embryos from Mahadevaiah et al. (*30*). **B)** Pseudotime (calculated using Slingshot, see **Materials and Methods**) of the injected cluster compared to E1.5 to E5.5 non-injected embryos. Dashed line delimits the E5.5-like, delayed and pre-EGA classification. **C)** X:A ratio heatmap per overlapping window (window size: 50 expressed genes, 49 genes overlap between consecutive windows) ordered by their X coordinate from 0 (top) to 87.5Mb (bottom). Single cells were clustered by k-means (k=5, see **Materials and Methods**). Top bars represent five clusters (cluster number in brackets), bottom bars indicate injected/non-injected status and developmental stage as identified in (**B**). The unedited cluster is split into non-injected cells and injected cells for visualisation. **D)** Box plots of genome coverage on the X in all identified clusters, normalised by the average in wildtype E5.5 cells. The X chromosome was divided into three regions: upstream of the *RSX* edit, the *RSX* expected edit (1700 bp) and downstream. The unedited cluster was divided into non-injected cells (E5.5) and injected cells (unedited (inj)). **E)** Heatmap of the ratio of representative single-cell genome reads at all the SNPs found in our colony for the different clusters, binned by 10 megabases along the X chromosome (see also **Materials and Methods**). **F)** Boxplot of *RSX* expression for E4.5 and E5.5 and all microinjected cells grouped by clusters. **G)** Box plot showing the X:A ratio of each single cell for E4.5 and E5.5 non-injected and injected cells grouped by clusters. (**F-G**) p-values calculated using a Wilcox unpaired test, outliers not shown. Unedited cluster is divided by non-injected cells (E5.5) and injected cells (unedited (inj)).

To reveal potential X-inactivation phenotypes in our injected cells, we devised a sliding X-to-autosome (X:A) ratio window approach. X:A ratios were calculated across successive, overlapping intervals along the X chromosome, to generate a nuanced, region-by-region measure of X-linked expression (**table S2**; see **Materials and Methods**). K-means clustering was then used to group cells into five clusters based on their X:A profiles across the X chromosome. Sliding X:A ratios for non-injected E4.5 cells were plotted alongside the E5.5 clusters for comparison (**Fig. 2C** and **fig. S2D**). For each cluster, we assessed genome coverage at the *RSX* target region (**Fig. 2D**), X-chromosome complement (**Fig. 2E**), *RSX* expression (**Fig. 2F**) and conventional, whole chromosome-based X:A ratio analysis to corroborate our sliding X:A ratio approach (**Fig. 2G**).

Within the largest cluster (cluster 1), the *RSX* target region was preserved, both X chromosomes were present, *RSX* was expressed, and the X:A ratio resembled that in non-injected E5.5 cells, as assessed by the novel sliding approach and by conventional whole chromosome X:A analysis (**Fig. 2C-G**). This cluster was therefore unedited.

The remaining four clusters exhibited CRISPR-induced mutations and aberrant X:A ratios. In cluster 2, the *RSX* target region was deleted, both Xs were preserved, and *RSX* expression was diminished, demonstrating successful *RSX* ablation. The X:A ratio was elevated relative to non-injected E4.5, E5.5 and injected unedited cells, linking *RSX* loss to increased X-expression (**Fig. 2C-G**). Cluster 2 was therefore *RSX* edited.

Cluster 3 carried a large deletion originating at a breakpoint within the first exon of *RSX*, which removed all downstream *RSX* exons and the entire 48.8 Mb distal region of the X chromosome. SNP analysis revealed that this deletion was on the paternal X, thus this cluster was named Xi arm loss. Accordingly, *RSX* expression was reduced, and the X:A ratio in the remaining centromeric interval was elevated, again demonstrating X-inactivation failure (**Fig. 2C-G**).

Cluster 4 also exhibited loss of heterozygosity, but on this occasion affecting the maternal X, hence this cluster was named Xa arm loss. *RSX* expression was unaffected, but the X:A ratio was unusually low, presumably because many genes on the active X were deleted (408 out of 702 expressed genes; **Fig. 2C-G**).

The final cluster (5; miscellaneous, misc) comprised a heterogeneous population of cells with mixed phenotypes that included partial X chromosome loss (**Fig. 2C-G**).

Our clustering revealed that most *RSX* edited, Xi arm loss and Xa arm loss cells fell in the delayed E4.5-like population (**Fig. 2C**; bottom row of heatmap, and **fig. S2D)**. This finding supported our earlier observation that X-dosage imbalance is associated with developmental delay (**fig. S1F**). Our results established two findings. Firstly, *RSX* is required for the initiation of X-inactivation as gleaned from sliding and whole-chromosome X:A ratio calculations. Secondly, CRISPR-Cas9 genome editing induces loss of heterozygosity in opossum embryos.

### *RSX* disruption causes biallelic X-expression

We next examined the effects of *RSX* deletion on the expression of specific X-genes. To account for the potential effect of developmental delay, we compared our *RSX*-edited cells to both the unedited cluster (injected and non-injected E5.5, age-matched) and E4.5 cells (non-injected, development-matched). Consistent with our X:A ratio findings, *RSX-*edited cells exhibited upregulation of genes across the X chromosome relative to control E5.5 cells (**Fig. 3A** and **fig. S3A)** and E4.5 cells **(fig. S3B** and **S3C**). We found a notable exception of a cluster of genes at the non-centromeric end (first three megabases) of the chromosome (**Fig. 3A** and **fig. S3B**). Genes upregulated in *RSX-*edited cells were strongly biased to the X chromosome relative to the autosomes (**Fig. 3B** and **fig. S3D**). SNP analysis revealed, as expected, maternally-biased expression of most X-genes in control E5.5 and E4.5 cells (median Xp:Xm expression ratio of 0.09 and 0.014, respectively; **Fig. 3C, table S3**). Exceptions included the known X-inactivation escapees *ATRX* and *CD99L2* (*23*), as well as potentially novel escapees or late-inactivating genes (**table S3**). However, in *RSX-*edited cells, X-gene expression was predominantly biallelic (median Xp:Xm (Xi:Xa) expression ratio of 0.52; **Fig. 3C**). The increased expression in *RSX*-edited cells affected genes that in wildtype cells were completely silenced (e.g., *VMA21* and *SMARCA1*) or partially silenced (e.g., *GLA* and *RLIM*; **Fig. 3D**). In conclusion, *RSX* deletion perturbs silencing of the paternal X in female opossum embryos.

**Fig 3.**
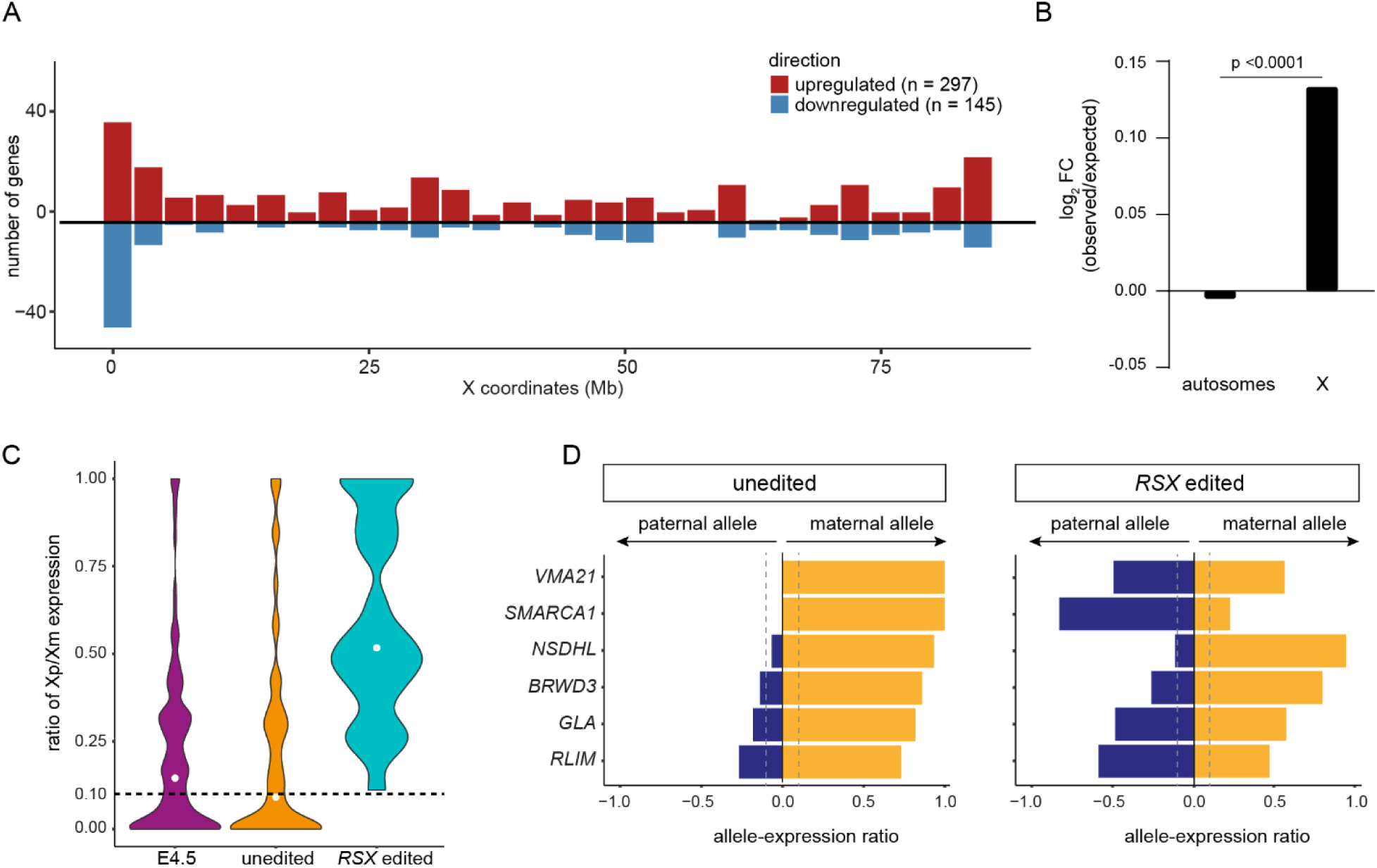
*RSX* deletion leads to increased expression from the paternal X at E5.5. **A)** Manhattan plot of the number of upregulated (red, n=297) and downregulated (blue, n= 145) genes per 3 Mb bins in the *RSX* edited cluster compared to the control. **B)** Bar plot of the ratio between the observed and expected log_2_ fold change in upregulated genes for the X and autosomes, in the *RSX* edited versus control comparison. P-value < 0.001, after Fisher’s exact test. **C)** Violin plot showing the allelic ratio of paternal (inactive) X (Xp) versus maternal (active) X (Xm) expression in E4.5, control and *RSX* edited clusters for genes where a SNP was present. A ratio of less than 0.1 is considered maternally expressed (monoallelic, dashed line). Median value is shown as a white dot. **D)** Bar plot showing the paternal and maternal allelic ratio in the control and *RSX* edited clusters, for six genes where a SNP was present, excluding known escapees (see also **Material and Methods**). A ratio of less than 0.1 from the paternal allele is considered monoallelic, maternally expressed.

### *RSX* has minor effects on maintenance of X-inactivation in opossum fibroblasts

When *Xist* is deleted in adulthood, some genes on the inactive X reactivate, but most remain silent (*14*). Maintenance of silencing may be mediated by CpG methylation, a highly stable epigenetic mark deposited at gene promoters on the eutherian inactive X (*32–34*). Interestingly, the marsupial inactive X lacks promoter DNA methylation, and is globally DNA hypomethylated relative to the active X (*23–27*). We therefore examined whether the absence of promoter DNA methylation renders the maintenance of marsupial X-inactivation more dependent on *RSX*.

To address this question, we deployed the same guide RNA pair used in embryos to generate three independent *RSX-*deletant immortalised female opossum fibroblast cell lines (**fig. S4A**). *MSN* expression, as determined by RNA-FISH, remained monoallelic in *RSX-*deletant fibroblasts (**Fig. 4A** and **fig. S4B**). Bulk RNA-seq revealed that *RSX* deletion did not overtly influence the X:A ratio or lead to a chromosome-wide increase in X-gene expression (**Fig. 4B-4C** and **fig. S4C-D**). We observed a subtle enrichment for upregulated genes on the X relative to the autosomes (**Fig. 4D**). SNP analysis confirmed that most X-genes remained monoallelic in *RSX* deletant cells, with some exceptions (13 out of 104 SNP-informative genes), including *RLIM*, *TIMM8A* and *HPRT1* (**Fig. 4E**, **fig. S4E**, and **table S4**). We then assayed genes that contained informative SNPs in both fibroblasts and the embryo. In all six cases, de-repression of the paternal allele was less pronounced in fibroblasts than in the embryo (**Fig. 3D and 4F**). We also observed increased expression of some X-inactivation escapees, including *PABIR2/3*, *PHF6* and *UTP14A* (**fig. S4E**). Increased expression of escapees on the inactive X has also been observed following *Xist* deletion in mouse cells (*15*, *35*).

**Fig 4.**
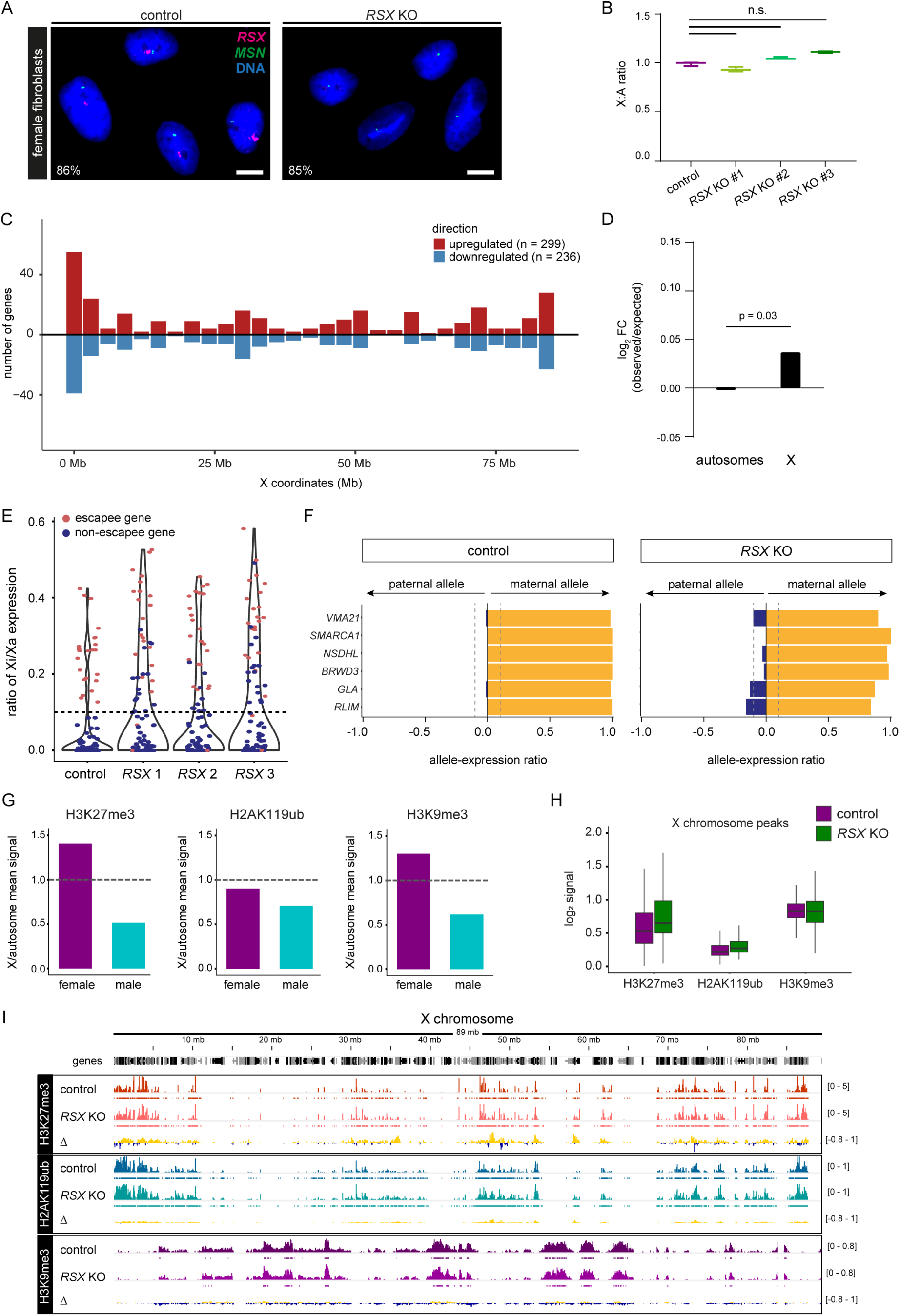
*RSX* deletion causes minor effects on the maintenance of X-inactivation. **A)** RNA-FISH of *RSX* (magenta) and *MSN* (green) in control and *RSX* KO female fibroblasts. Nuclei were stained with DAPI. Scale bar, 10 µm. **B)** Box plot of X:A ratio in control and *RSX* KO fibroblast lines. n.s. = no significant difference (p-value > 0.05 after Kruskal-Wallis test). **C)** Manhattan plot of the number of upregulated (red) and downregulated (blue) genes per 3 Mb bins in the *RSX* KO versus control opossum fibroblasts. **D)** Bar plot of the log_2_ fold change between the number of observed and expected upregulated genes in the X and autosomes in the *RSX* KO versus control comparison. P-value = 0.03, after Fisher’s exact test. **E)** Violin plot showing the allelic ratio of inactive X (Xi) versus active X (Xa) expression ratio in the different fibroblast lines. Escapee genes (allelic ratio >0.1 in controls) are marked in red. **F)** Bar plot showing the paternal and maternal allelic ratio in control and *RSX* KO fibroblasts, for six genes where a SNP was found in our colony, excluding known escapees (see also Fig. 3D). A ratio of less than 0.1 from the paternal allele is considered monoallelic, maternally expressed. **G)** Bar plot showing the ratio between the mean CUT&Tag signal on the X peaks and the mean signal on the autosome peaks for H3K27me3, H2AK119ub and H3K9me3. **H)** Box plot of the log_2_ signal of the three different histone marks in all the X chromosome peaks identified in female control and *RSX* KO fibroblasts. **I)** profile of the three histone marks along the X chromosome in control (top) and *RSX* KO (middle). Bottom track shows the difference (Δ) between both tracks.

Our findings demonstrated that *RSX* deletion caused more minor effects on the maintenance compared to the initiation of X-inactivation. Given that DNA methylation does not regulate X-inactivation in marsupials, we profiled three other repressive modifications implicated in eutherian X-inactivation using CUT&Tag, and inferred Xi patterns by comparing profiles in females versus males (**fig. S5A**). H3K27me3, deposited by the Polycomb Repressive Complex 2 (PRC2), and the heterochromatin mark H3K9me3, are enriched on the opossum inactive X (*20*, *21*, *23*), but H2AK119ub, catalysed by the Polycomb Repressive Complex 1 (PRC1), has to our knowledge never been examined in marsupials.

At the genome-wide level, H3K27me3 and H2AK119ub marks were enriched at promoters and intergenic regions, while H3K9me3 was predominantly present at intergenic regions, as predicted (**fig. S5B**). H3K27me3 and H3K9me3 were enriched on the X chromosome in female compared to male fibroblasts (**Fig. 4G** and **fig. S5C-S5D**). Unexpectedly however, the X-enrichment for H2AK119ub typical of eutherians (*36–39*) was not observed in the opossum, suggesting that this mark may not contribute to X-inactivation in marsupials (**Fig. 4G** and **fig. S5D**). The profile of all three marks was overtly unaffected in *RSX* KO cells (**Fig. 4H-4I** and **fig. S5E**). We conclude that *RSX* is not required to maintain H3K27me3 and H3K9me3 on the marsupial inactive X. Furthermore, deletion of *RSX*, like *Xist,* has a greater impact on the initiation than the maintenance of X-inactivation.

## Discussion

The discovery of *RSX* was published in 2012, but resolving its function in marsupial X-inactivation has been hampered by the technical complications of genome editing in marsupials. Here, we overcome this challenge, demonstrating that *RSX* is required for the initiation of X-inactivation. Our findings highlight the remarkable evolutionary flexibility in dosage compensation mechanisms, wherein *RSX* and *Xist* can drive X-inactivation despite exhibiting no primary sequence homology.

Deletion of *RSX,* similarly to *Xist,* causes more modest effects on the maintenance compared with the initiation of X-silencing, at least as gleaned from our *in vitro* fibroblast system. H2AK119ub is dispensable for the maintenance of X-inactivation in the mouse embryo proper (*38*) and, based on our findings, it may likewise be dispensable in the opossum. Moreover, while promoter DNA methylation is enriched on the eutherian inactive X, marsupials retain X-inactivation in the absence of this modification. These observations suggest that some modifications associated with X-inactivation may be more dispensable for maintenance than previously thought, and raise the possibility that conserved yet undiscovered epigenetic mechanisms regulate X-inactivation maintenance in both eutherians and marsupials.

Our study also has important implications for the development of marsupial genome editing. Marsupials are deployed in diverse research areas, including neuroscience, cancer biology and antibiotic research. We demonstrate that while CRISPR-Cas9 genome editing is feasible, it also induces large chromosome deletions, a phenomenon also observed in mouse and human embryos (*41*, *42*). These findings highlight both the promise and the technical challenges of applying genome editing to marsupials, and emphasise the importance of comprehensively characterising edited alleles as comparative functional genomics in these fascinating species continues to expand.

## Acknowledgements

We would like to thank the people and teams that helped us during this project: The Biological Research Facility and veterinarians of the Francis Crick Institute for maintaining the opossum colony; Agustina Resasco, Agata Zielinska, Sarah Phelan and Katherine Courtis for their help with embryo transfer in London; Yuliia Dovga for her technical help; the Vertebrate Genomes Project, in particular Erich Jarvis, Giulio Formenti, Bonhwang Koo and Jen Balacco for generating the mMonDom1 opossum assembly; Jasmin Zohren and Christopher Barrington for their help in preparing a custom opossum genome assembly; Jeremie Subrini for providing a plasmid for genome editing; Steven Henikoff and Vangelis Christodoulou (Crick Structural Biology platform) for providing pA-Tn5 enzyme for CUT&Tag; Akira Konishi for his scientific drawings; the Francis Crick Institute Genomics, Scientific Computing, and Advanced Light Microscopy science technology platforms for their assistance; Edith Heard, members of her lab, and members of the J.M.A.T. laboratory for discussion and feedback.

## Author contributions

J.M.A.T. conceived the project. A.C., S.M., S.O., D.S., H.K. and J.M.A.T. designed the experiments. S.O. and T.A. designed the CRISPR-Cas9 approach. A.C. and K.I. performed opossum zygote microinjection. S.W., R.Y. and A.S. performed embryo transfer. A.C. performed RNA FISH in embryos. A.C. and S.M. recovered embryos for single-cell omics. D.S. designed the scG&T-seq. W.V. and A.C. performed computational analysis in embryos. S.O. and S.M. generated *RSX* KO fibroblast lines and performed RNA FISH. D.S. and S.M. characterised *RSX* deletions. F.D. generated immortalised fibroblast lines and performed initial tests. S.M. performed and analysed bulk RNA-seq and CUT&Tag in opossum fibroblasts. J.L.V provided additional opossums. S.M., A.C. and J.M.A.T wrote the manuscript with input from all authors.

## Data availability

Sequencing data generated in this study have been deposited in the National Center for Biotechnology Information’s Gene Expression Omnibus (GEO) database (accession numbers pending) and Sequence Read Archive (SRA) database (accession numbers pending).

## Supplementary Figures

**Figure S1.**
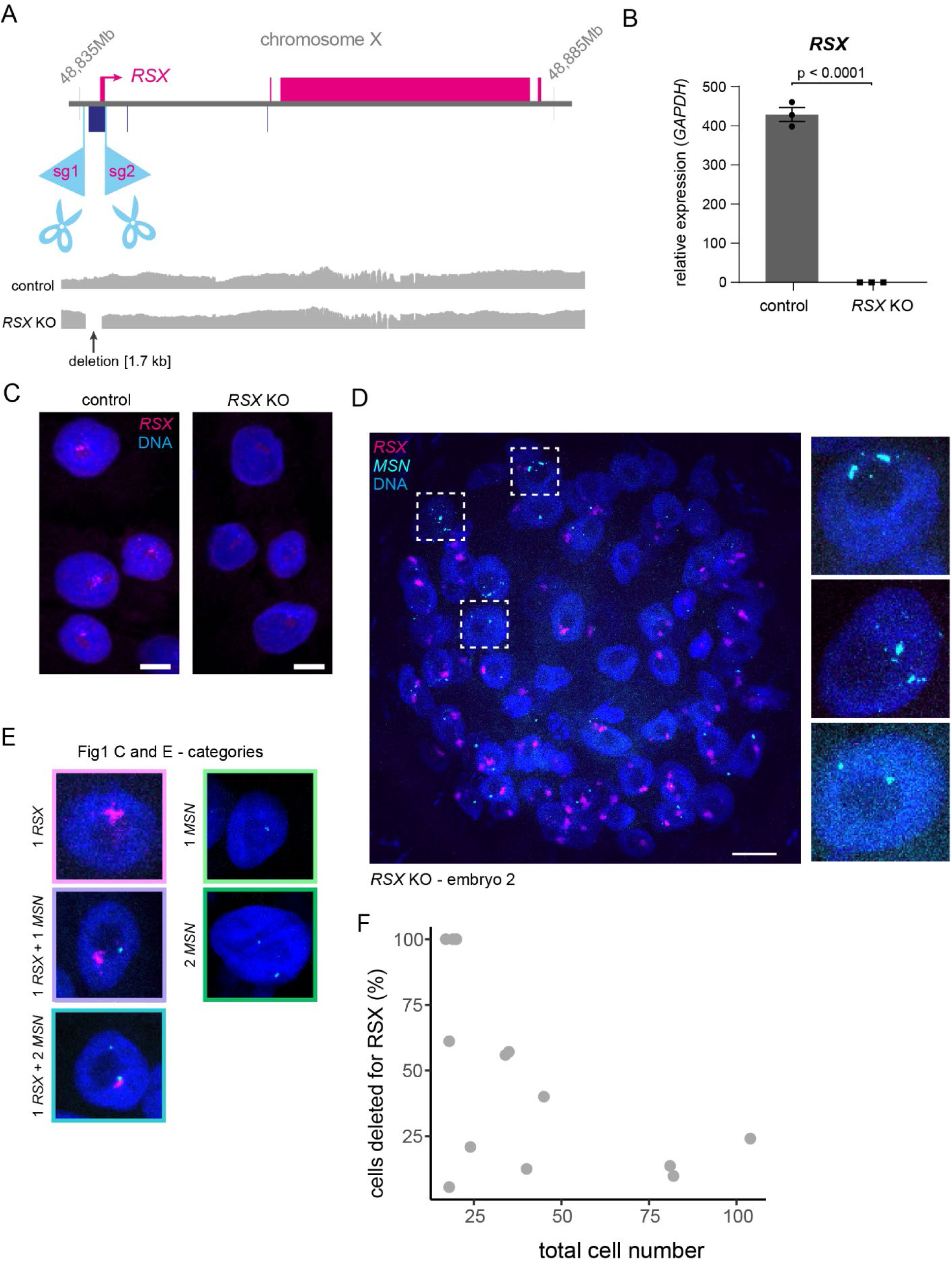
*RSX* deletion in the opossum. **A)** Schematic representation of the *RSX* locus showing the targeted area with single-guide RNAs (sg1-2). A fibroblast *RSX* KO cell line shows the expected deletion (∼1.7 kb) in the targeted area, which includes the first *RSX* exon, as identified with nanopore targeted sequencing. **B)** qPCR results showing that RSX expression is abrogated in *RSX* KO cell line (p-value < 0.001 by Student t-test). **C)** RNA FISH for *RSX* (magenta) in control and *RSX* KO cells. Nuclei were stained with DAPI. Scale bar, 10 µM. **D)** Confocal Z-projection of female opossum embryo microinjected with guides targeting *RSX* (embryo 2), with RNA FISH for *RSX* (magenta) and *MSN* (cyan). Scale bar 20µm. Insets shows three cells magnified. **E)** Representative cells of each of the categories from **Fig. 1C and E**. **F)** Scatter plot of the total cell number versus the percentage of cells deleted for *RSX* for all the embryos classified as female in our RNA FISH analysis.

**Figure S2.**
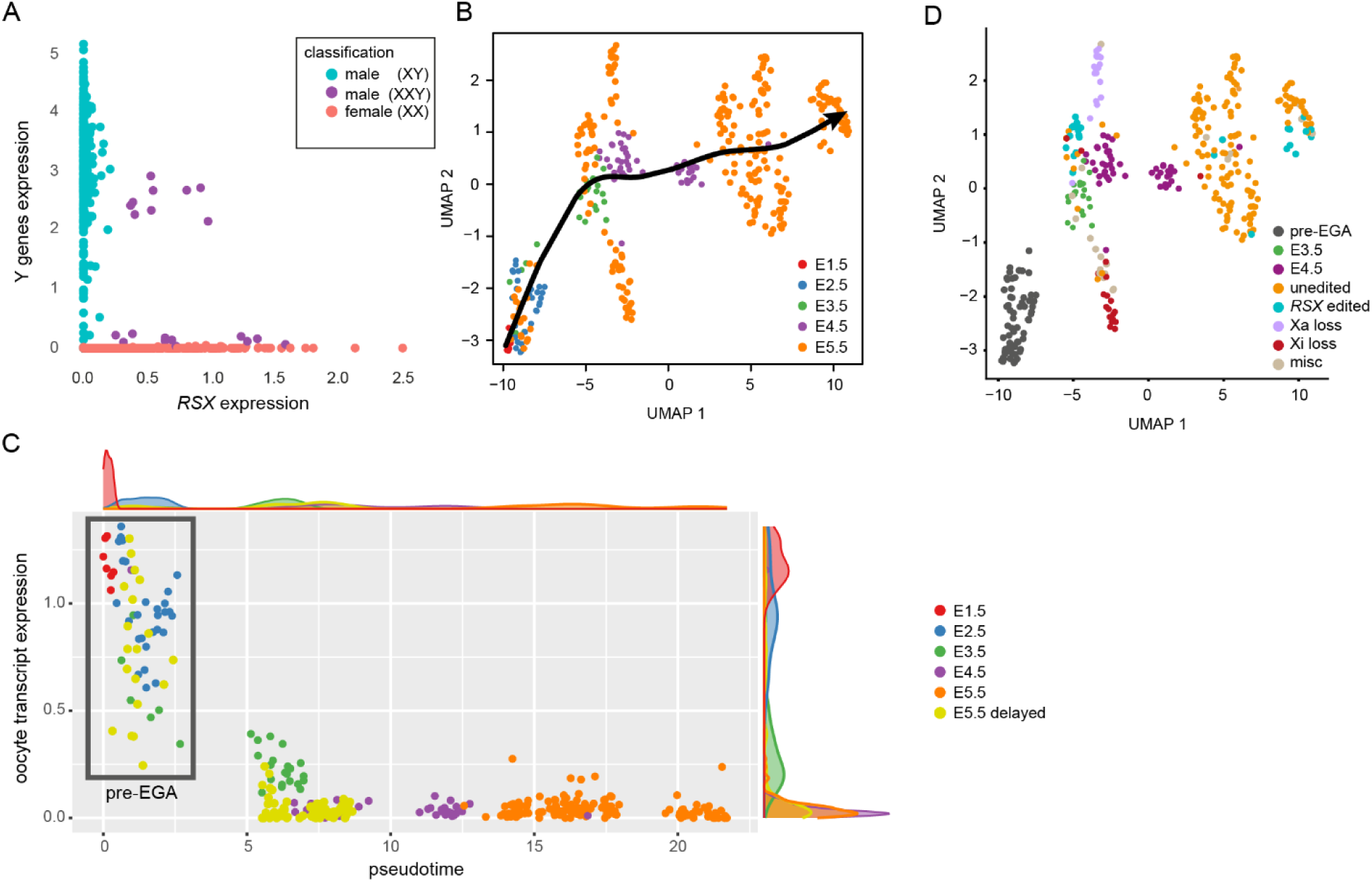
Identifying the *RSX* edited population in our single-cell analysis. **A)** Scatter plot of *RSX* expression versus Y gene expression in each single cell. Y gene expression was defined as the sum of all the genes expressed from the first Y scaffold NW_026605000.1. Some embryos appeared to be of XXY and were excluded. **B)** UMAP representation of single cells from all the XX E5.5 injected and from E4.5 and E5.5 non-injected embryos, as well as single cells from E1.5, 2.5 and 3.5 non-injected embryos from Mahadevaiah et al. (*30*). The Slingshot pseudotime trajectory is shown as a black arrow. **C)** Scatter plot of the Slingshot pseudotime and oocyte transcript expression (see also **table S1**) of each single cell coloured by developmental time (E1.5 to 5.5) or pseudotime (delayed E5.5) with density ridges on the side. A pseudotime of 5 was defined as the threshold for embryonic genome activation (EGA). **D)** Same UMAP as **(B)** coloured by X:A clusters.

**Figure S3.**
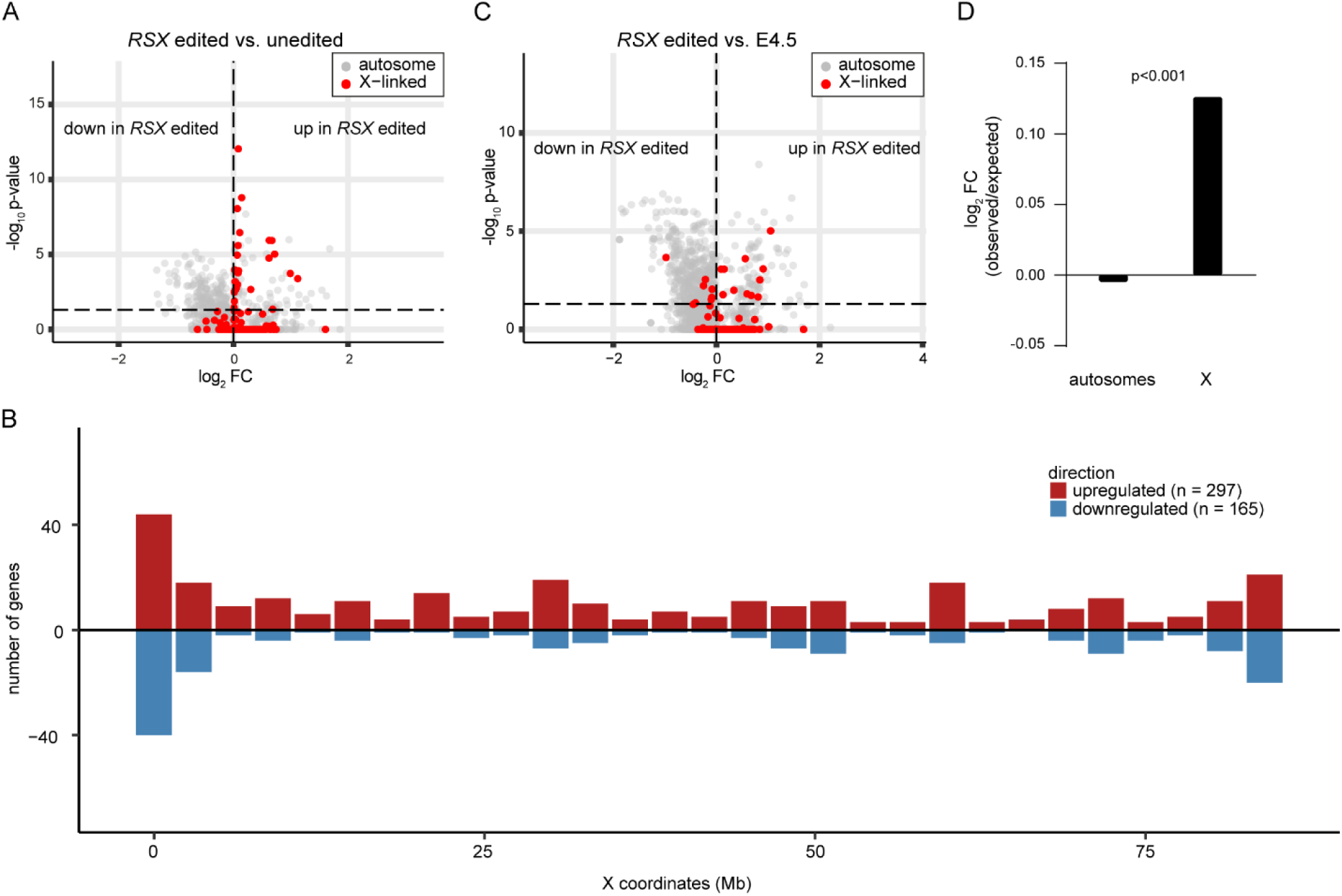
Comparison of *RSX* edited versus the E4.5 cluster. **A)** Volcano plot showing differential gene expression between *RSX* edited versus the unedited (E5.5) cluster. X-linked genes are marked in red. **B)** Manhattan plot of the number of genes per 3Mb bins upregulated (red, n=297) and downregulated (blue, n= 165) in the *RSX-*edited cluster compared to non-injected E4.5. **C)** Volcano plot showing differential gene expression between *RSX* edited and non-injected E4.5 cells. **D)** Bar plot of the ratio between the observed and expected log_2_ fold change in upregulated genes for the X and autosomes, in the *RSX* edited versus non-injected E4.5 comparison. P-value < 0.001 after Fisher’s exact test.

**Figure S4.**
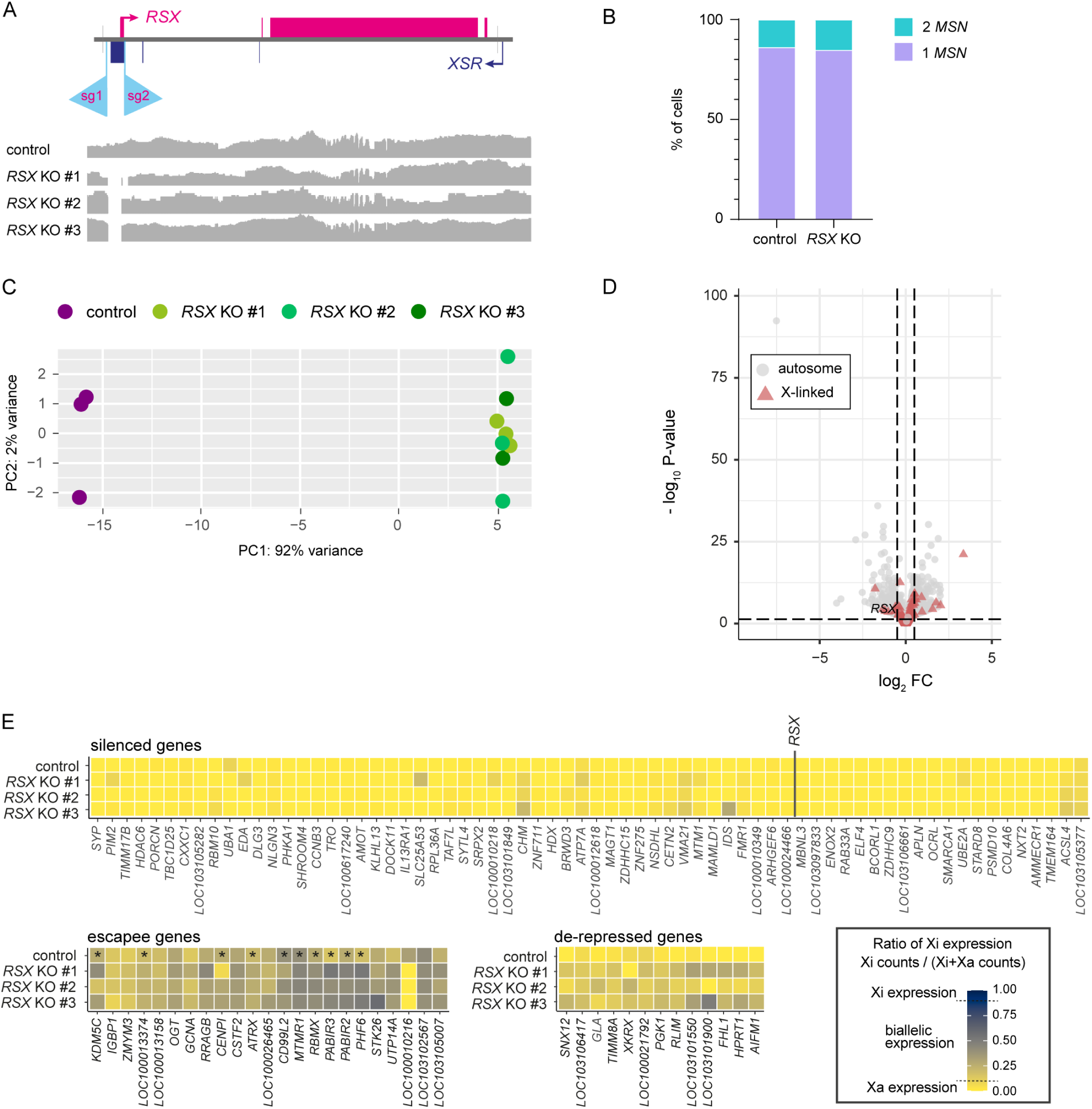
Role of *RSX* in X-inactivation maintenance. **A)** Schematic representation of *RSX* locus showing targeted area with single-guide RNAs (sg1-2). Three fibroblast *RSX* KO cell lines show the expected deletion (∼1.7 kb) in the targeted area, which includes the first *RSX* exon, as identified with nanopore targeted sequencing. **B)** Percentage of fibroblasts expressing one or two *MSN* alleles by RNA-FISH in control (n=130) and *RSX* KO (n=138) cells. **C)** Principal Component Analysis (PCA) of bulk RNA-seq of the three *RSX* KO clones and one control line. The first component (PC1) separates the samples by genotype. **D)** Volcano plot showing differentially expressed genes (DEG) in *RSX* KO lines versus control. X-genes are represented by red triangles, while autosomal genes are shown in grey circles. **E)** Heatmaps showing the average allele-specific expression ratio of X-linked genes in control and three *RSX* KO female fibroblast clones. The 104 genes were selected based on the presence of informative SNPs. Genes, ordered by genomic position, are distributed in three categories: silenced genes (ratio of Xi expression < 0.1 in control and at least two *RSX* KO clones), escapee genes (ratio of Xi expression > 0.1 in control) and de-repressed genes (ratio of Xi expression > 0.1 in at least two *RSX* KO clones). Genes marked with * were previously identified escapee genes (*23*).

**Figure S5.**
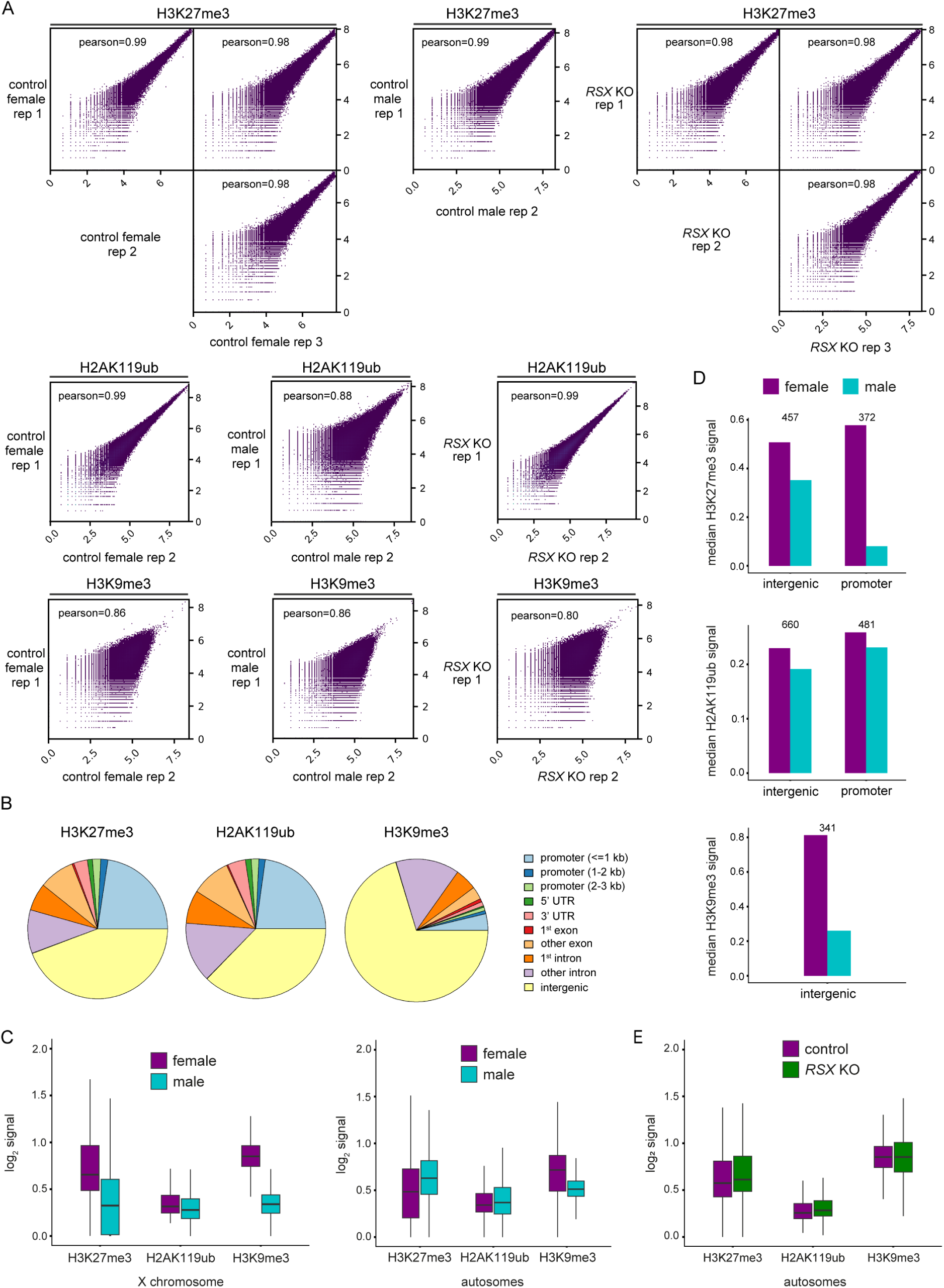
Repressive histone marks in the opossum. **A)** Scattered plots showing correlation of CUT&Tag signal for H3K27me3, H2AK119ub and H3K9me3 in control female, control male, and *RSX* KO replicates. Pearson correlation coefficient is shown for each comparison. **B)** Pie chart representing the proportion of peaks for each genomic feature for H3K27me3, H2AK119ub and H3K9me3 in female fibroblasts. **C)** Box plots of the log_2_ signal of three different histone marks in all the peaks identified in the X chromosome (left panel) and autosomes (right panel) of male and female fibroblasts. **D)** Bar plots showing the median CUT&Tag signal for H3K27me3 (top plot), H2AK119ub (middle plot) and H3K9me3 (bottom plot) in peaks identified on the X chromosome of female and male opossum fibroblasts associated with intergenic and promoter (<1 Kb from transcription start site) regions. The number of peaks included in the analysis is annotated at the top of each graph. **E)** Box plot of the log_2_ signal of the three different histone marks in all the peaks identified in the autosomes of control female and *RSX* KO fibroblasts.

## Materials and Methods

### Animal welfare

Opossums were maintained in the Francis Crick Institute Biological Research Facility in accordance with UK Animal Scientific Procedures Act 1986 regulations (project licence P8ECF28D9) and subject to Francis Crick Institute’s internal ethical review. No randomization or blinding was performed, and no statistical methods were used to predetermine sample size. Opossums were individually housed in double-decker cages (GR1800, Tecniplast), with male and female animals housed in separate rooms except during mating periods. The temperature of the housing was maintained between 24 °C and 28 °C, and humidity was maintained between 55% and 75%, with a 14 h/10 h light/dark cycle (21h-7h). Opossums had free access to dried food and water, supplemented every second day with live mealworms, and weekly by fresh fruit. To induce oestrous before mating, adult male and female (≥6 months) opossums were placed in a single-storey rat cage with a central delimiter for 2 days and then swapped into each other’s side for an additional 2 days. Subsequently, delimiter was removed, and they were kept together for up to 7 days, during which period animals were monitored by infrared CCTV cameras for mating behaviour (*45*). The animal experiments conducted at RIKEN Kobe Campus were approved by the Institutional Animal Care and Use Committee of RIKEN Kobe Campus (Approval No. A2001-03). Animal housing at RIKEN Kobe Campus was set as previously described (*31*).

### Preparation of a modified MonDom1 reference genome and annotations

mMonDom1 (GCF_027887165.1; GCF_027887165.1_mMonDom1.pri_genomic.fna) assembled by the Vertebrate Genomes Project (https://vertebrategenomesproject.org/)(46) and the Oz Mammals Genomics (https://ozmammalsgenomics.com/), was downloaded from NCBI and used with the following modifications. Coordinates for *RSX* and *XSR* coordinates were added to the reference gtf file (*RSX.1*, *XSR.1*, GCF_027887165.1_mMonDom1.pri_genomic), based on Mahesh Sangrithi analysis (*30*). An additional transcript for *XSR* was added (*XSR.2*), based on trinity as well as an additional transcript for *UBE1Y*, which we called *UBE1Y-like* on the Y chromosome, and that seemed to encompass the end of the *UBE1Y* transcript that wasn’t captured in the official genome annotation. In addition, a mitochondrial chromosome from MonDom5 Ensembl 99 release was added to the fasta and the gtf file, allowing for single-cell quality checks and filtering. Lastly, we renamed the LOC NCBI ids from all the gametologs on the Y chromosome that were described previously and of *NONO* on the X chromosome. To ensure compatibility with nf-core cutandrun pipeline, the modified mMonDom1 genome was split into smaller fragments of ∼100 Mb for CUT&Tag analysis. For all modifications made to mMonDom1, see **table S5**.

### CRISPR design

Single-guide RNAs (sgRNAs) were designed using CRISPR direct (crispr.dbcls.jp/). sgRNAs were annealed and ligated into the targeting plasmid px458 (Addgene plasmid #48138) after adding a second cloning backbone for sgRNA. The px458 plasmid has Cas9 from *S. pyogenes* with 2A-EGFP. The guides result in a 1711 bp deletion encompassing the putative promoter region and most of the first exon of *RSX*. RSX-sg1: GCTGCCTCTTCTGACGCTAA tgg. RSX-sg2: AATGACAGGGACCATCCGAG agg. A mutation was present in some individual in the London colony. When needed, a different guide was used to target the mutated allele. RSX-sg2-mut: AAGGACAGGGACCATCCGAG agg. An extra pair of guides were tested in Japan that resulted in the same outcome by RNA FISH. RSX-sg3: GTCACTGTCAGCAGACCGGA tgg. RSX-sg4: TGTAGCCTCACTACAATACC tgg.

### Embryo genome editing

In Kobe (Japan), everything was performed as described previously (*31*). In London (UK), embryos were collected 28 hours after coitum or at 2:00 (in the morning), whichever came first (average mating time is between 21:00 and 22:00 in our colony; dark cycle from 21:00 to 7:00), in FHM (Hepes buffered medium). Embryos were microinjected using a water microinjection system (Eppendorf CellTram Vario) controlled by a micromanipulator (Narishige MMO4). Capillaries were pulled using a puller (Sutter instrument – P97). Drilling was performed thanks to mercury loaded in the tip of the microinjection capillary using a PrimeTech piezo unit (PMAS-CT150). Embryos were held by a holding needle and positioned so that one or both pronuclei were visible. When possible, the CRISPR mix (crRNAs (IDT) at 50 ng/μL after annealing with tracrRNA (IDT), HiFi Cas9 (IDT) 100 ng/μL in injection buffer) was injected into both pronuclei. Alternatively, single pronucleus injection or cytoplasmic microinjection was performed. Embryos were then placed in pre-equilibrated KSOM and left to recover in the incubator at 32.6°C, 5% CO_2_ until the embryo transfer (average time of the embryo transfer was 10:30). Before the embryo transfer, embryos were briefly washed in FHM. Embryos were inserted into the uterus of a pseudopregnant female as described previously (*31*). After inserting the embryos in the uterus, the uterus was stitched closed, which improved embryo recovery efficiency.

### Fibroblast genome editing

Primary opossum fibroblasts were derived from a newborn female animal and immortalized using SV40-tag virus infection. Opossum fibroblasts were maintained in DMEM (Gibco) supplemented with 20% fetal bovine serum, 1% GlutaMax (Gibco), 1% sodium pyruvate (Gibco) and 1% penicillin–streptomycin (Gibco, 10,000 U/ml).

Immortalized opossum fibroblasts were seeded onto gelatin-coated wells of 6-well plates. The following day, fibroblasts were transfected using PEI MAX (49553-93-7). 2 µg of plasmid (containing sgRNAs to target *RSX* or with no sgRNAs as control) was added to 200 µl of Opti-MEM (Gibco) and 8 µl of PEI MAX (1 mg/ml). After 48 hours, cells were dissociated, and GFP-positive single cells were picked under a stereomicroscope with a fluorescent lamp. Picked single cells were plated onto gelatin-coated wells of 96-well plates. Clones that proliferated and expanded were PCR-genotyped using the following primers flanking the targeted region: RSXdel_Fw: 5’-CTCTTCCGGTGTGGTGTCTC-3’; RSXdel_Rv: 5’- CACGTACTCACAACCCTGCT-3’. Three different lines with an *RSX* deletion and one control line were established. Further characterisation of the deletions was performed using Nanopore targeted sequencing to enrich the coverage on the X chromosome: High-molecular-weight DNA was extracted from fibroblast cell pellets using QIAGEN g-tips (QIAGEN), quantified by fluorescence using Qubit HS assay (Thermo Fisher Scientific). Approximately 5 µg of DNA was prepared for sequencing using SQK-LSK109 (Oxford Nanopore Technologies), following the manufacturer’s protocol. >500ng library was loaded onto a MinION R9 flow cell (Oxford Nanopore Technologies), with reloading every 16 hours. Adaptive sampling was run using a custom implementation of Readfish (*47*), selecting for ∼10MB of chrX around the *RSX* locus. Basecalling was performed using guppy v3.2.10; Analysis of nanopore data was performed using nf-core nanoseq (v. 1.1.0) (*48*) with the following options: --protocol DNA --aligner minimap2, and .bam files were visualised using IGV (v2.19.7).

### Embryo collection

Embryos were collected four and a half (E4.5) or five and a half (E5.5) days after coitum, between 10h-12h. In our colony, coitum happens in most of the cases between 21h and 22h. Uteri were cut out and washed briefly in warm (32°C) PBS. Uteri were cut open along the blood vessel and spread out in a new warm PBS-PVP (0.01% PVP in PBS) dish. Embryos were collected using a 600µm tip on a stripper pipettor and kept on ice in PBS- PVP.

### Embryo RNA FISH

E5.5 embryos were fixed in 4% PFA on ice for 30 min. They were then washed three times in PBS 0.1% triton X100 for 5 min, and permeabilised in methanol at -20°C, at least overnight. Embryos were then rehydrated and transferred to the hybridisation solution, incubated 30 min at 37°C and then transferred to the hybridisation solution containing the probes (*RSX* HCR v3., molecular instruments, (*49*)) and *MSN* (488 dUTP labelled BAC VM18-259O2 (CHORI)) (*30*) and incubated at 37°C for a minimum of 16 hours. Embryos were then washed and amplified using HCR hairpins (647nm), stained with Hoechst 33342 and mounted in VECTASHIELD PLUS (H-1900, vector laboratories). Embryos were imaged on a confocal microscope (Zeiss LSM 880, slice thickness 0.5µm).

### Embryo DNA FISH

DNA FISH was performed in injected embryos after RNA FISH and imaging to sex them. Embryos were recovered one by one and post-fixed for 30min in cold 4% PFA. They were then washed twice and DNA was denatured at 70°C for 7 min in 2x SSC-PVP and 14 min in 70% formamide in 2x SSC-PVP. Embryos were then cooled down for 5 min in -20°C methanol. They were then pre-hybridised for 30 min at 37°C in 50% formamide 50% 2x hybridisation buffer (4 x SSC, 50% dextran sulphate, 10mg/ml BSA, 2mM Vanadyl Ribonucleoside), and hybridised in the same solution containing *MSN* probes (647 dUTP labelled BAC VM18-259O2 (*50*) (CHORI)) and Y chromosome paint (Gold 525 Enzo dUTP labelled BAC MAB-198O, gift from D. Page (*51*)) for a minimum of 16 hours. Embryos were then washed, stained with Hoechst 33342 and mounted in VECTASHIELD PLUS (H-1900, vector laboratories). Embryos were imaged on a confocal microscope (Zeiss LSM 880, slice thickness 0.5µm).

### Embryo single-cell collection

Each embryo was processed individually as follows: The embryo shell was punctured using a needle and then the embryo was incubated in 5 mg/ml pronase (P8811-100MG, Sigma) for 5 min to get rid of the mucoid layer and the zona pellucida. The embryo was then transferred into a cold PBS-PVP drop and was dissociated using ∼20-30 µm opening glass capillaries. Single cells were collected using a 75 µm tip on a stripper pipettor and put in individual wells containing 2.5µL RLT+ buffer (Qiagen). Once collected, plates were frozen on dry ice and stored at -70°C. We collected all E4.5 and E5.5 embryos used in this paper.

### Single-cell G&T sequencing

Single-cell genome and transcriptome sequencing (scG&T-seq) was performed as previously described (*52*, *53*), with some minor modifications. Briefly, cells lysed in RLT+ buffer (Qiagen) were thawed and supplemented with biotinylated oligo-dT conjugated to MyOne streptavidin-coated magnetic beads (Thermo Scientific). Following incubation on a thermomixer at 2000 rpm for 20 min at room temperature, plates were transferred to a Biomek i7 liquid handling platform (Beckman Coulter Inc.), where supernatant was collected, and beads were washed twice with genomic DNA (gDNA) wash buffer. Oligo-dT-bound transcripts were eluted into FLASH-seq RT-PCR mix, which was the cycled 18 times to amplify cDNA (*54*). Genomic DNA-containing supernatant was cleaned with AMPure XP beads (Beckman Coulter Inc.), and whole-genome amplification performed following the manufacturer’s protocol at half-volume using Resolve DNA v1 (Bioskryb Inc.). Both cDNA and gDNA libraries were prepared using Nextera XT DNA kit (Illumina), and sequenced on either NovaSeq 6000 or NovaSeq X+ platforms using PE100 or PE150 configuration.

### SNP discovery in opossum parents

Parent WGS data was processed using nf-core (v24.10.2) sarek (v3.5.1). Reads were aligned using BWA against our custom opossum genome. Duplicates were marked using GATK and the resulting cram file was converted to a bam file. Aligned reads from each parent pair were jointly processed using bcftools (v1.19) to identify genomic variants relative to the reference genome (mpileup followed by call). Multi-allelic variants were split into separate records (norm), and only biallelic SNPs differing between the parents were retained for downstream analysis (view). The final filtered variant set was indexed (tabix) and converted into a tab-delimited table containing chromosome, position, reference and alternative alleles, genotype, read depth, and allele depth information (query) for downstream analysis. In R, a master SNP list was generated by collecting all unique SNPs across the six pairs of parents, in total 124,503 potential SNPs were identified. This master list was used to identify the presence of the active and inactive X in the offspring. Firstly, the parental master SNP list was filtered by minimum reads for mum (≥ 8 reads, bam-readcount) and dad (≥ 3 reads, bam-readcount) at each SNP location, after removing the duplicates (samtool, markdup) identified by the nf-core pipeline. Each SNP was considered further if all the reads were the same within one parent (homo/hemizygous) and different between each parent. For the transcriptome analysis, only the 49,854 SNPs from the list residing within 1kb of a gene were considered. In total, 93,424 SNPs were present in at least one pair of parents in the genome analysis, and 38,629 SNPs in the transcriptome analysis.

### Allelic ratio analysis

For the genome analysis, reads were extracted for each single cell based on their parents of origin. Reads were found in 19,031 SNPs across all single cells. SNPs were then binned per 10 megabases, only bins with at least two different SNPs were kept.

For the transcriptome analysis, reads were extracted for each single-cell and kept if there were at least 6 reads in total per embryo within a cluster for a particular SNP (626 informative SNPs). Subsequently, SNPs that belong to non-protein-coding genes were removed from the analysis (537 informative SNPs). Reads were then binned per gene (173 informative genes).

### Single-cell transcriptomic analysis

Using nf-core (v25.10.0) RNAseq (3.14.0) the raw read data were aligned using star_rsem to the modified MonDom1.pri genome. Low-quality cells were filtered out using the SingleCellExperiment R package (1.20.0) using the following thresholds. Cells with a high mitochondrial % (30.7%, 3 MADs), low gene coverage (2000 genes), high rRNA % (9.73%, 5 MADs) or low read count (1980, bottom 5%) were excluded from the analysis, leaving 635 high-quality cells. The counts data were then processed in Seurat (4.3.0). Lowly expressed genes were filtered out using the function CreateSeuratObject() and min.cells=3. The data was further processed using the NormalizeData(), FindVariableFeatures(), ScaleData() and RunPCA() functions. To integrate all batches together, the Seurat object was split by batch using SplitObject(), followed by NormalizeData() and FindVariableFeatures() for each batch. Integration was done using SelectIntegrationFeatures(), FindIntegrationAnchors(), then IntegrateData(), specifying all remaining genes as the ‘features.to.integrate’ parameter and k.weight=30. Cells were genotyped using an *RSX* vs sumY plot, which identified XX, XY and XXY cells. The XXY embryo was excluded, and the remaining cells were classified into XX and XY using Y expression, with cells with ≤ 0.1 sum Y normalised expression classified as XX. To be confident in our XX cell classification, embryos which had a mix of XX and XY cells were classified as XY. In total, 295 cells were genotyped as XX and retained for further analysis. Next, we created an XX reference dataset by integrating only the E4.5 and E5.5 WT XX cells from this study (67 cells) with WT E1.5, E2.5 and E3.5 XX cells (63 cells) from a previous study (*30*) using SelectIntegrationFeatures(), FindIntegrationAnchors() then IntegrateData(). The WT XX UMAP was generated by using ScaleData(), RunPCA(), FindNeighbors() then RunUMAP(). The E5.5 microinjected XX cells (228 cells) were then integrated into the WT XX UMAP, again using SelectIntegrationFeatures(), FindIntegrationAnchors(), and then IntegrateData(). This created the final UMAP containing 358 XX cells. To infer a pseudotime trajectory, we used the R package slingshot (2.10.0), using the function slingshot(), setting the starting cluster to E1.5, specifying approx_points=250. This generated one pseudotime trajectory. Using the E1.5 and E2.5 cells as reference, together with pseudotime values and a list of maternally enriched genes, we identified 20 E5.5 cells in our dataset, which we classified as pre-EGA. The remaining E5.5 cells were then classified using our sliding X-to-A ratios method (see later). DEGs were identified using FindMarkers, specifying logfc.threshold=0.00001) and an adjusted p-value of 0.05. Volcano plots were generated using the R package EnhancedVolcano (1.16.0). Manhattan barplots were generated by binning the X chromosome into 3Mb bins and calculating the number of genes that had a positive or negative log2 fold change. Over-representation of upregulated X-linked genes was assessed by calculating the log_2_ fold change between the observed and expected numbers of upregulated genes on the X chromosome relative to the autosomes. Upregulated genes were defined as genes with a positive log_2_ fold change (>0) in the transcriptional analysis. The number of expected genes was calculated under the assumption that upregulated genes were distributed in proportion to the total number of expressed genes on each chromosome. Statistical analysis was performed using Fisher’s exact test.

### X- to-autosome clustering of E5.5

Sliding X:A ratios were performed on all E4.5 and E5.5 cells apart from those E5.5 cells classified as pre-EGA. Candidate autosome and X-linked genes for the X:A calculation were identified by selecting genes that were expressed ≥0.1 normalised counts in at least 70% of cells (192 / 275 cells). This identified 4,805 autosomal genes and 95 X-linked genes. To implement the sliding approach, candidate X-linked genes were ordered by coordinate, then grouped into windows of 50 genes, with a 49 gene overlap, thereby creating 46 windows spanning the entire X chromosome. For each window, the mean autosomal and X-linked expression per cell was calculated using colMeans(). The X:A ratio per cell was calculated by dividing the chrX mean by autosomal mean. In order to identify potential *RSX* edits, we focused on the E5.5 cells and applied kmeans clustering, with k=5. Set.seed() was defined as 1 to ensure reproducibility. The X-to-A ratios and clusters were visualised as a heatmap using the R package ComplexHeatmap (2.14.0), revealing five groups of E5.5 cells, each representing a different edit.

### Fibroblast RNA FISH

Fibroblasts grown on slide chambers were washed with ice-cold PBS and permeabilised with 0.5% Triton in PBS for 10 minutes on ice. Cells were then fixed with 4% PFA in PBS for 10 minutes on ice. After washed, fibroblasts were dehydrated through ice-cold 70%, 80%, 95% and 100% ethanol for 3 min each, and air-dried. BAC VM-18-303M7 (CHORI) was used for *RSX* RNA FISH, and BAC VM-18-25902 (CHORI) was used for *MSN* RNA FISH. BAC DNA was labelled using Nick Translation Kit (Abbott) with fluorescent nucleotides (spectrum orange-dUTP, 02N33-050; spectrum green-dUTP, 02N32-050, Abott), and cells were hybridized with a denatured mix of probes along with 1 µg salmon sperm DNA in hybridization buffer (50% formamide, 10% dextran sulfate, 1 mg/ml PVP, 0.05% Triton X-100, 0.5 mg/ml BSA, 1 mM vanadyl ribonucleoside complex in 2× SSC) at 37 °C overnight in a humid chamber. The slides were then washed in a coplin jar three times for 5 min in 50% formamide in 1× SSC preheated to 45 °C, and three times for 5 min in 2× SSC preheated to 45 °C. Samples were mounted on Vectashield containing DAPI (Vector laboratories) with a coverslip. Imaging was performed in an Olympus SoRa Spinning disc, and images were then visualised and analysed in Fiji (*55*).

### Fibroblast RNA purification and qRT–PCR

RNA from control and *RSX* KO female opossum fibroblasts was purified using RNAqueous-Micro Total RNA Isolation kit (Invitrogen, AM1931). Purified RNA was reverse-transcribed using the Maxima First Strand cDNA synthesis kit (Thermo Scientific, K1641). Loss of *RSX* expression was confirmed by qRT-PCR. The following primers were used to assess gene expression in qRT–PCR assays using PowerUp SYBR Green (Applied Biosystems, A25780): RSX (5′-AGAAGGGACCCCAAGACAC-3′, 5′-TGGGTCACTTCCACTTCCTC-3′); GAPDH (5′-TAAATGGGGAGATGCTGGAG-3′, 5′-ATGCCGAAGTTGTCGTGAA-3′).

### Fibroblast bulk RNA-sequencing

Libraries from control and *RSX* KO RNA samples were prepared with the NEBNext Ultra II Directional PolyA mRNA kit according to the manufacturer’s instructions. Libraries were sequenced on the Illumina NovaSeq 6000 system (paired end, 100-bp read length). Raw RNA-seq reads were processed using the RNA-seq nf-core pipeline (v3.2); star_rsem was used to generate raw read counts. The read counts were processed in R using the DESeq2 package (v1.36). Genes expressed at very low levels were filtered out by applying a rowSums filter of ≥5 to the raw counts table. Raw counts were normalized using the DESeq() function, specifying ∼genotype in the design formula. log2[fold change] and adjusted P values between *RSX* KO and control samples were calculated using the lfcShrink() function in DESeq2, specifying type = ‘ashr’. PCA was calculated using plotPCA with default parameters (using top 1000 more variable genes).

X:A ratios were calculated using the median expression of X and autosomal genes in each sample after filtering out genes expressed at low levels (transcripts per million (TPM) of <1). X:A ratios between control and *RSX* KO samples were compared by Kruskal-Wallis test.

Over-representation of upregulated X-linked genes was assessed by calculating the log_2_ fold change between the observed and expected numbers of upregulated genes on the X chromosome relative to the autosomes. Upregulated genes were defined as genes with a positive log_2_ fold change (>0) in the DESeq2 analysis. The number of expected genes was calculated under the assumption that upregulated genes were distributed in proportion to the total number of expressed genes on each chromosome. Statistical analysis was performed by Fisher’s exact test.

### Fibroblast CUT&Tag

CUT&Tag was performed in opossum fibroblasts as previously described (*56*). Control female, control male and the RSX KO #3 clone were used. Approximately 200,000 cells per sample were used. After washing in wash buffer (20 mM Hepes at pH 7.5, 150 mM KCl, 0.5 mM spermidine, and 1× protease inhibitors), cells in each sample were incubated with 10 µL of concanavalin A-coated magnetic beads (Bangs Laboratories, BP531) for 10 min. Bead-bound cells were resuspended in dig-wash buffer (wash buffer and 0.05% digitonin) containing 2 mM EDTA, 0.1% BSA and a 1:100 dilution of the corresponding primary antibody (rabbit anti-H3K27me3, Cell Signaling #9733; rabbit anti-H2AK119ub1, Cell Signaling #8240; rabbit anti-H3K9me3, abcam #8898). Primary antibody incubation was performed overnight at 4°C, followed by incubation with secondary antibody (Guinea Pig anti-Rabbit antibody, ABIN101961; 1:100 dilution in dig-wash buffer) for 1 hour at room temperature (RT). Samples were then washed twice with dig-wash buffer and incubated with pA-Tn5 adapter complex, using a 1:250 dilution in dig-300 buffer (0.05% digitonin, 20 mM Hepes at pH 7.5, 300 mM KCl, 0.5 mM spermidine, and 1× protease inhibitors) for 1 hour at RT. After washing twice in dig-300 buffer, cells were then resuspended in 100 μl of tagmentation buffer (dig-300 buffer with 10 mM MgCl_2_) and incubated at 37°C for 1 hour. To stop fragmentation and solubilize DNA fragments, 20 mM EDTA, 0.1% SDS and 0.2 mg/ml Proteinase K were added to each sample and incubated at 50°C for 1 hour. DNA was subsequently extracted via phenol-chloroform isoamyl alcohol for library preparation using phase-lock tubes (Qiagen MaXtract High Density #129046). DNA libraries were prepared using NEBNext Ultra II Q5 master mix (NEB #M0544S) mixed with a universal i5 and a uniquely barcoded i7 primer (*57*). Libraries were purified using 0.9x AMPure XP beads (Beckman Coulter), subjected to tapestation analysis, and Illumina sequencing. Samples incubated with non-specific IgG antibody (rabbit anti-IgG, abcam #ab46540) were used as negative controls, but not sequenced because the low background of the technique resulted in negligible libraries.

The pA-Tn5 adapter complex was prepared as described (see protocol at https://www.protocols.io/view/3xflag-patn5-protein-purification-and-meds-loading-j8nlke4e5l5r/v1).

### Fibroblast CUT&Tag analysis

Data was processed using the nf-core cutandrun pipeline (*48*, *58*) (https://nf-co.re/cutandrun/) with the following options: -r 3.1 --normalisation_mode CPM --use_control false --peakcaller SEACR --extend_fragments true --replicate_threshold 2. Correlation analyses were performed using multiBamSummary and plotCorrelation from deepTools (v3.5.1) (*59*). Peak annotation was performed in female control fibroblasts using ChIPseeker (*60*, *61*). To compare the CUT&Tag signal in female versus male, and female control versus *RSX* KO samples, we generated a master peak file with all the consensus peaks identified for each mark using bedtools merge (*62*). We generated one master peak list for each paired comparison. The signal for each histone modification was then quantified in those peaks using multiBigwigSummary (deepTools) and plotted using ggplot2.

### SNP analysis in fibroblasts

In female fibroblasts, SNPs on the X chromosome were identified using the genomic Nanopore targeted-sequencing data (see Fibroblast genome editing section). FreeBayes with a threshold (-C) of 20 reads was used to identify SNPs. Allele-expression ratios based on SNPs in gene loci were then assessed in the bulk RNA-seq data of the control and three *RSX* KO clones. A total of 232 SNPs within 104 expressed X-genes were found (**table S4**).

